# Upd-like/JAK/STAT signaling promotes regeneration and tolerance to bacterial infection in the *Aedes* midgut

**DOI:** 10.1101/2025.09.21.677592

**Authors:** P.H. Vieira, B. Hixson, J. Hodgson, N. Buchon

## Abstract

*Aedes aegypti*, a major vector of arboviruses, relies on its midgut as a key interface with microbial communities that influence vector competence. While ingestion of pathogenic bacteria is known to trigger both immune and regenerative responses in the mosquito gut, the signaling pathways that govern epithelial renewal remain poorly defined. Here, we address this gap by performing transcriptomic profiling of the *Aedes aegypti* midgut following oral infection with entomopathogenic bacteria. We uncover a broad transcriptional activation of immune pathways, including IMD, Toll, and notably, the JAK-STAT signaling cascade. Functional knockdown experiments reveal that JAK-STAT signaling is essential for infection-induced epithelial cell proliferation and survival, but dispensable for bacterial clearance, supporting a specialized role in tissue repair and infection tolerance. We further identify a previously uncharacterized cytokine-like gene, *Upd-like*, structurally related to Drosophila Unpaired proteins and mammalian cytokines, which is strongly induced upon bacterial challenge and activates JAK-STAT signaling in mosquito cells. Silencing *Upd-like* markedly impairs epithelial regeneration after infection, establishing it as a key regulator of gut homeostasis. Together, these findings demonstrate a conserved function for JAK-STAT signaling in midgut epithelial renewal and identify Upd-like as a novel cytokine mediating regenerative responses during bacterial infection.

## Introduction

Mosquito-borne diseases remain a major global health challenge, with *Aedes aegypti* serving as the principal vector for arboviruses such as dengue, Zika, and chikungunya^1–3^. The mosquito midgut is a critical interface between the insect and its environment, functioning both as a site of nutrient absorption and as a frontline barrier against pathogens encountered through blood or sugar feeding^4,5^. Beyond digestion, the midgut harbors a resident microbiota and regulates pathogen access to systemic tissues, thus shaping the mosquito’s vector competence^4,6–9^. Recent studies have shown that the midgut epithelium comprises multiple cell types, including absorptive enterocytes (ECs), hormone-secreting enteroendocrine cells (EEs), and basal progenitors resembling intestinal stem cells (ISCs)^8,10^. The variety of specialized cells in the midgut allows it to adjust its functions and structure in response to different challenges. As in Drosophila, epithelial turnover occurs via regulated ISC proliferation and differentiation, a process triggered by feeding, microbial exposure, or tissue damage, which is essential for maintaining barrier integrity^11–15^. Although accelerated epithelial renewal has been documented following enteric infection and chemical injury^16–18^, the molecular programs orchestrating epithelial regeneration in mosquitoes, and their conservation relative to Drosophila, remain largely unknown.

In addition to its structural complexity, the mosquito gut is immunologically active, constantly interacting with microbes, whether symbiotic, opportunistic, or pathogenic^8,9,19^. To manage this microbial exposure, *Aedes aegypti* relies on conserved innate immune pathways, primarily Toll, and IMD, which coordinate transcriptional responses against a broad spectrum of pathogens. These pathways regulate the expression of antimicrobial peptides (AMPs), lysozymes, and other effector enzymes through the activation of NF-κB transcription factors like Rel1 and Rel2 ^20–23^. Such responses constitute classic resistance mechanisms, in which the host seeks to eliminate or limit pathogen proliferation^24^. However, resistance is only part of the equation: growing evidence from both mosquito and Drosophila studies supports a complementary strategy known as tolerance, defined as the ability to limit infection-induced damage without necessarily decreasing pathogen load^24–28^. In this light, tissue turnover, particularly ISC activation and epithelial repair, could be viewed as a form of tolerance that sustains gut structure and function during microbial stress. Yet, it remains unclear whether mosquitoes employ such tolerance strategies in the gut. Moreover, how they balance resistance and tolerance, particularly in the context of midgut physiology, microbiota interactions, and long-term effects on vector competence remains poorly understood.

In *Drosophila melanogaster*, the integration of immune defense and epithelial regeneration has been well characterized. The midgut epithelium also includes polyploid ECs, EEs, and undifferentiated progenitors (ISCs and enteroblasts, EBs), which dynamically interact to preserve tissue integrity and function^11,12,15,29^. ISCs adjust their proliferation, differentiation, and endocycling rates in response to nutrient levels, hormonal cues, microbiota composition, pathogens, oxidative stress, and aging^11–13,30,31^. This process is mediated by a complex network of immune and developmental pathways, including JAK-STAT, EGFR, and Wingless, that link EC stress to ISC proliferation via niche reprogramming^11–13,32,33^ . Despite the recognized importance of midgut cell dynamics in mosquitoes, it remains unclear whether such extensive reprogramming of midgut signaling occurs in response to bacterial ingestion, and if so, what genetic networks drive epithelial renewal.

The JAK-STAT pathway is a fundamental and evolutionarily conserved signaling cascade in insects, playing critical roles in both immune responses and developmental processes^22,30^. Activation begins when extracellular cytokine-like ligands bind to specific cell surface receptors, triggering Janus kinases (JAKs) to phosphorylate Signal Transducers and Activators of Transcription (STATs). Phosphorylated STATs then dimerize and translocate to the nucleus, where they regulate the expression of immune-responsive genes^34^ . Negative regulators such as Suppressor of Cytokine Signaling (SOCS), provide feedback inhibition and serve as transcriptional readouts of pathway activity^35^. In Drosophila, the JAK-STAT pathway is a key regulator of both immunity and epithelial regeneration^30,36,37^. It is activated by the Unpaired (Upd) family of ligands, Upd1, Upd2, and Upd3, with Upd3 being strongly induced by bacterial infection, linking this pathway to epithelial damage and infection-driven responses^14,36–39^. In *Aedes aegypti*, JAK-STAT acts as a key antiviral mechanism, restricting replication of viruses such as dengue, Zika, and West Nile independently of other immune pathways^22,40–42^. These findings position JAK-STAT as a key regulator of mosquito immunity, however, its roles in antibacterial immunity, gut epithelial renewal and tolerance to bacterial infection remain largely unexplored^22,40,43^. Furthermore, the cytokine-like ligands that activate JAK-STAT signaling in the mosquito midgut remain largely undefined, as no clear Unpaired (Upd) homologs have been identified in *Aedes* genomes. In the context of viral infection, the secreted peptide Vago (VLG-1) has been proposed as a cytokine-like activator of the JAK-STAT pathway. Upon viral challenge, Vago is upregulated in a Dicer-2–dependent manner and secreted, leading to JAK-STAT activation and restriction of viral replication, as shown in Culex mosquitoes infected with West Nile virus^41,44,45^. This antiviral mechanism is functionally analogous to the mammalian interferon response, positioning Vago as an IFN-like cytokine^41^. Notably, Vago appears to signal through a receptor distinct from the canonical JAK-STAT receptor Domeless, suggesting a noncanonical activation route^22^. In contrast, the Upd family in Drosophila represents canonical JAK-STAT ligands that act through Domeless to regulate immunity, regeneration, and homeostasis^38^. While both Vago and Upd can activate the pathway, they are induced by different stimuli and likely engage distinct signaling mechanisms. Although Vago’s role in antiviral defense is well established, its function in gut physiology remains unclear, and additional ligands in mosquitoes have yet to be identified.

Here, to investigate how *Aedes aegypti* responds to ingested bacteria, we combined transcriptomic profiling, functional genetics, and cellular assays to systematically dissect midgut immune and regenerative responses. Our goal was to identify master regulators of resistance and tolerance mechanisms that maintain gut homeostasis and may ultimately influence vector competence. We examined the response to two Gram-negative pathogens in both laboratory-adapted and field-derived mosquito strains, uncovering a conserved transcriptional program involving immune, developmental, and stress-related pathways. We found that the JAK-STAT pathway is induced upon bacterial challenge and is essential for ISC proliferation and epithelial renewal. Moreover, we identified and characterized a previously unrecognized Unpaired-like cytokine that activates JAK-STAT signaling and is required for epithelial renewal and host survival. Together, our findings uncover fundamental mechanisms governing gut regeneration and immune homeostasis in mosquitoes, and highlight key molecular targets for manipulating tolerance, resistance, and ultimately vector competence.

## Results

### Oral Bacterial Infection Stimulates Immune, Stress and Developmental Processes in the *Aedes aegypti* Midgut

To investigate cellular and molecular responses to enteric bacterial infection in *Aedes aegypti*, we performed RNA sequencing (RNA-seq) on midguts from two mosquito strains: a field-derived Thai strain, representing a genetically diverse population with phenotypes reflective of natural conditions, and the laboratory-adapted Liverpool (Lvp) strain, an isogenic line widely used for mechanistic studies. This dual approach allowed us to capture both natural variability and reproducible responses under controlled laboratory settings.

Under unchallenged conditions, transcriptome profiling revealed distinct strain-specific expression programs (Supplementary Fig. 1a). Volcano plots highlighted numerous transcripts elevated in either Thai (blue) or Lvp (red) backgrounds. Gene Ontology (GO) enrichment showed that Thai-upregulated genes were enriched for translation, protein targeting, spliceosomal assembly, and metabolic pathways (Supplementary Fig. 1b). By contrast, Lvp-enriched genes were associated with protein phosphorylation, signal transduction, receptor activity, and cellular organization (Supplementary Fig. 1c). These findings underscore fundamental molecular differences between the two strains at baseline.

Female mosquitoes were orally challenged with the Gram-negative bacteria, *Erwinia carotovora carotovora* (*Ecc15*), a non-lethal pathogen that robustly activates gut immunity and epithelial renewal, and *Pseudomonas entomophila* (*Pe*), a highly virulent bacterium that causes severe gut damage. These species were selected as well-characterized models for dissecting mosquito responses to naturally encountered enteric infections and tissue repair. Following infection, midguts were collected for RNA-seq to profile transcriptional changes associated with immune activation and tissue regeneration. For Lvp, sampling was performed at 12 hours post-infection (hpi) with *Ecc15*, whereas Thai midguts were profiled at 4, 12, and 24 hpi following infection with either *Ecc15* or *Pe*. Principal component analysis (PCA) of transcriptomic profiles revealed clear stratification of samples by genotype (PC1) and treatment (PC2, Fig. 1a). PC2 captured infection progression, with untreated controls clustering at the top of the plot, while Ecc15- and Pe-infected samples formed distinct groups at the bottom. Although the two mosquito genotypes displayed different basal expression patterns (PC1), they followed similar trajectories upon infection, indicating broadly comparable responses to gut pathogens. Together, these findings demonstrate that oral bacterial infection profoundly reprograms gene expression in the mosquito midgut.

**Figure 1.**
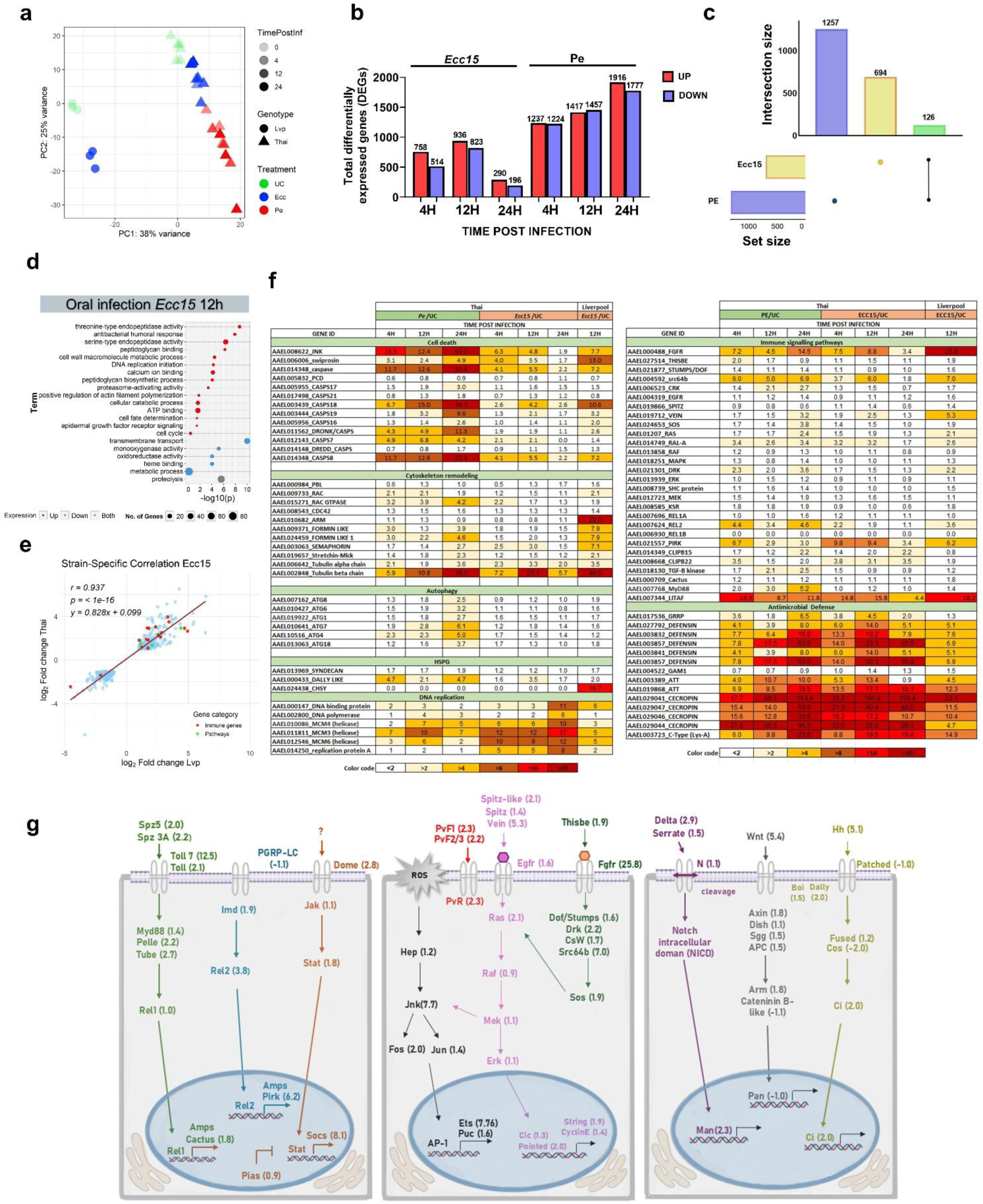
Transcriptional landscape and immune signaling dynamics in *Aedes aegypti* midgut upon oral bacterial infection. (a) Principal component analysis (PCA) of mosquito midgut transcriptomes. PC1 (38%) separates genotypes (Lvp, circles; Thai, triangles), while PC2 (25%) reflects infection progression. Colors indicate treatments (UC, green; Ecc15, blue; Pe, red), and shading denotes time post-infection (0–24 h, light to dark). (b) Number of differentially expressed genes (DEGs) following *Ecc15* and *Pe* infection compared to sugar-fed controls at 4–24 h post-infection. Red bars indicate upregulated and blue bars downregulated genes. (c) Overlap of DEGs between infections, highlighting infection-specific and shared transcriptional responses. (d) Gene Ontology (GO) enrichment analysis of DEGs at 12 h post-Ecc15 infection, showing enriched biological processes. Circle size reflects the number of genes per GO term, and color indicates expression direction. (e) Strain-specific fold-change correlation of DEGs in Ecc15 infection, comparing Thai and Lvp strains. Red line indicates regression, dashed line indicates y = x. (f) Heatmaps showing expression of selected immune and signaling pathway genes across conditions. (g) Schematic of signaling pathways (Imd, Jak-STAT, EGFR, Notch, Hedgehog) transcriptionally modulated during infection with ECC15 in Liverpool. fold changes at 12 hpi are shown in parentheses. All experiments were performed in three biological replicates.

Differential expression analysis of the Thai strain transcriptome confirmed that *Pe* elicited a more robust and sustained transcriptional program than *Ecc15*, reflected in a larger number of differentially expressed genes (DEGs) over time (Fig. 1b). While the infections shared a core DEG set of 126 genes, *Pe* induced a broader and more persistent midgut response (Fig. 1c).

As early as 4 hpi, both *Ecc15* and *Pe* triggered widespread transcriptional changes in Thai midguts involving hundreds of genes (Fig. 1b; Supplementary Fig. 2a, d). Shared DEGs were enriched for proteolytic enzymes (serine proteases, metalloproteinases, proteasome subunits) and canonical immune effectors such as cecropins, lysozymes, and peptidoglycan recognition proteins. In parallel, genes encoding metabolic transporters (e.g., aquaporins, trehalose transporters) were downregulated, suggesting metabolic remodeling of the gut environment (Supplementary Fig. 2g).

At 12 hpi, transcriptional signatures diverged between infections. *Ecc15* strongly induced immune response genes alongside those linked to epithelial growth, mitosis, and tissue regeneration (Fig. 1d). In contrast, *Pe* activated pathways associated with cellular stress, apoptosis, and actin cytoskeleton remodeling, indicating greater cytotoxicity (Supplementary Fig. 2h). Patterns of downregulation also differed: *Ecc15* transiently repressed transmembrane transport and actin polymerization genes, possibly to limit tissue damage, whereas *Pe* broadly suppressed vacuolar transport, proteolysis, and regeneration pathways, consistent with more severe disruption of midgut homeostasis.

To assess strain variation in the response to enteric pathogens, we compared *Ecc15*-induced expression changes across our two mosquito strains. Pearson correlation analysis showed strong concordance (r = 0.937), indicating highly conserved transcriptional responses between Thai and Lvp mosquitoes. Key immune and signaling genes were regulated in the same direction, underscoring robustness of core midgut responses despite genetic background differences.

Infections strongly upregulated antimicrobial peptides, including defensins, attacins, and cecropins (Fig. 1f), which are essential for microbial control and maintaining microbiota balance. Stress-associated genes linked to apoptosis, autophagy and cytoskeletal remodeling were also induced, likely facilitating the removal of damaged cells and subsequent tissue reorganization. At the same time, activation of DNA replication-related genes pointed to increased intestinal stem cell proliferation, suggesting an acceleration of epithelial renewal (Fig. 1f). Although the magnitude of regulation varied between bacterial species and mosquito strains, the overall transcriptional trends were consistent, reflecting a conserved midgut response to enteric infection.

Analysis of signaling components revealed a dynamic, time- and strain-dependent induction of multiple signaling pathways. Genes from the receptor tyrosine kinase (RTK), mitogen-activated protein kinase (MAPK), Toll, and immune deficiency (IMD)pathways were broadly upregulated, with expression typically peaking at 12 hpi, especially following *Ecc15* infection (Fig. 1f).In the Lvp strain, *Ecc15* strongly induced RTK pathway members (Fgfr, Egfr, Spitz, Ras, Raf, Mek2) along with regulators of stress and regeneration including Jnk, Notch, Wnt, and Hedgehog. This finding supports a model in which immune and regenerative programs are tightly coordinated in response to bacterial challenge.

To integrate these findings, we assembled DEGs into a schematic representation of signaling pathways at 12 hpi with *Ecc15* in Lvp (Fig. 1g). This model illustrates coordinated activation of immune (Toll, IMD, JAK-STAT), stress (JNK), and developmental (RTK, Notch, Wnt, Hedgehog) networks. Together, our results reveal a complex transcriptional program that integrates pathogen recognition, immune defense, and epithelial renewal. These data demonstrate that *Aedes aegypti* mounts a robust and multifaceted midgut response to enteric bacterial infection, built upon conserved signaling cascades that jointly regulate immunity and tissue homeostasis.

### JAK-STAT Pathway controls tolerance to infection

To investigate the role of the JAK-STAT pathway (Fig. 2a) during bacterial infection in *Aedes aegypti*, we analyzed gene expression following oral challenge with *Ecc15* or *Pe*. Transcriptomic data revealed significant upregulation of core JAK-STAT pathway components in both the Liverpool and Thai mosquito strains, indicating a conserved activation of this signaling cascade in response to bacterial exposure (Fig. 2b). RT-qPCR validation confirmed strong induction of STAT, which showed the highest fold change among all pathway components at 12 hpi (Figs. 2c,e). The receptor Domeless was also upregulated, further supporting pathway activation. While expression of the negative regulator PIAS (protein inhibitor of activated STAT) remained relatively unchanged, *Socs*, a known transcriptional target and inhibitor of JAK-STAT signaling, was robustly induced by both infections in both strains, serving as a clear marker of pathway activation (Figs. 2d,f). Together, these results demonstrate that enteric bacterial infection elicits a strong and conserved activation of the JAK-STAT pathway in the mosquito midgut.

**Figure 2.**
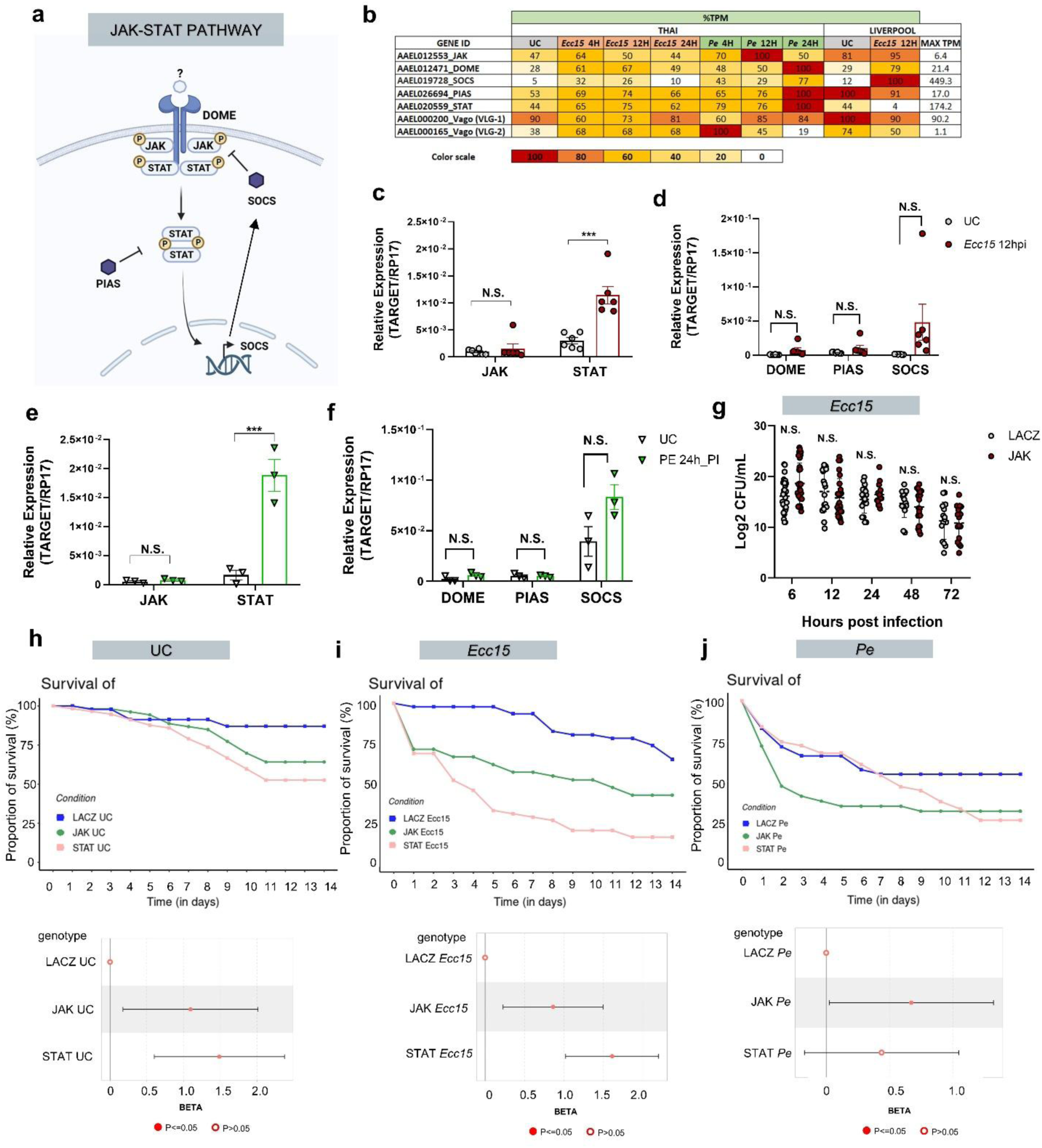
JAK-STAT signaling is activated by oral infection and promotes host survival without affecting bacterial load. (a) Schematic representation of the JAK-STAT pathway in *Aedes aegypti*, showing canonical components: receptor DOME, kinases JAK, transcription factor STAT, and regulators SOCS and PIAS. (b) Heatmap of transcript levels (FC) for Jak-STAT pathway components in midguts of mosquitoes after *Ecc15* and *Pe* oral infection, across time points and strains. (c-f) RT-qPCR validation of STAT target gene induction, confirming infection-induced JAK/STAT activation in response to *Ecc15* (c-d) and *Pe* (e-f). Relative expression data were analyzed using an unpaired Student’s t-test to compare control and treated groups for each gene. Results are presented as mean ± SE. Each dot represents one biological replicate (n=20). Statistically significant differences are indicated by asterisks (\**p* < 0.05), while “N.S.” denotes no significant difference. (g) Bacterial load quantification in mosquitoes injected with dsRNA targeting JAK or control (LacZ), followed by *Ecc15* infection, showing no significant difference in bacterial CFU over time. (h-j) Survival analysis of mosquitoes silenced for JAK or STAT, under sucrose (h), *Ecc15* (i), or *Pe* (j) infection. JAK-STAT silencing significantly reduces survival in infected groups, suggesting a protective role during gut infection. Insets: hazard ratio plots from Cox proportional hazards models.

To elucidate the functional consequences of JAK-STAT activation during bacterial infection, we performed RNA interference-mediated knockdown of key pathway components, including STAT and JAK (Supplementary Fig. 3a, c). Despite robust transcriptional activation of the pathway, knockdown of these genes did not significantly alter bacterial load compared to control mosquitoes, suggesting that JAK-STAT signaling does not directly mediate bacterial clearance in this context (Fig. 2g). However, survival analyses revealed that STAT-silenced mosquitoes exhibited significantly reduced survival following infection with both *Ecc15* and *Pe*, as well as under uninfected conditions (Figs. 2h–j). Survival analysis further confirmed that STAT knockdown increased mortality, underscoring the pathway’s critical role in promoting host survival during bacterial challenge. These findings indicate that the JAK-STAT pathway contributes primarily to immune homeostasis or tolerance, rather than direct antimicrobial activity, by mitigating infection-induced physiological stress and supporting tissue integrity.

In summary, our results demonstrate that the JAK-STAT pathway is transcriptionally activated in response to bacterial infection in *Aedes aegypti*, with STAT acting as a key effector. While not essential for bacterial clearance, the pathway is crucial for maintaining mosquito survival during infection, likely through modulation of infection tolerance and preservation of gut homeostasis. These findings expand the known roles of JAK-STAT signaling in mosquitoes, highlighting its importance beyond antiviral defense.

### JAK-STAT signaling is essential for infection-induced midgut intestinal stem cell proliferation and epithelial renewal

Given its established role in tissue repair in Drosophila, JAK-STAT signaling may contribute to host tolerance against bacterial infection by promoting epithelial regeneration^26,28,46^. Previous studies, including our own, have shown that oral infection with pathogenic bacteria such as *Ecc15*, *Serratia marcescens*, and others triggers a strong proliferative response in the mosquito midgut epithelium. This is evidenced by increased numbers of phospho-histone H3 (PH3)-positive cells and enhanced nucleotide analog incorporation, reflecting elevated mitotic activity and DNA synthesis following infection or chemical damage (e.g., paraquat, SDS)^16–18^. We thus hypothesized that the JAK-STAT pathway supports host survival by coordinating this regenerative response.

To test this, we first confirmed that bacterial infection induces epithelial renewal and established the time course of the response. Midgut proliferation was assessed by quantifying DNA synthesis (EdU incorporation) and mitosis (PH3 immunostaining) following oral infection with *Ecc15* or *Pe*. Confocal imaging revealed minimal EdU incorporation in uninfected (UC) controls, indicating low baseline proliferation (Supplementary Fig. 4a). Following *Ecc15* infection, EdU-positive cells progressively increased through 72 hpi. Quantification confirmed a significant rise in EdU+ cell percentages over time relative to controls Supplementary Fig. 4b). Similarly, numbers of PH3+ mitotic cells per gut increased sharply, peaking at 72 hpi (Supplementary Fig. 4c). A comparable response was observed after *Pe* infection: EdU incorporation and PH3+ cell numbers were both significantly elevated compared to uninfected controls (Supplementary Fig. 4d–f). These findings demonstrate that enteric bacterial infection induces robust and sustained epithelial proliferation in the mosquito midgut, detected as early as 12 hpi.

To determine whether JAK-STAT signaling is required for this proliferative response, we performed RNAi-mediated knockdown of JAK and STAT, followed by infection and quantification of epithelial proliferation. In *Ecc15*-infected controls (dsLacZ-injected), we observed a strong increase in EdU+ cells and PH3+ mitotic figures (Fig. 3a– c). However, knockdown of either JAK or STAT resulted in a near-complete loss of EdU incorporation and PH3 staining, with both metrics reduced to baseline levels comparable to uninfected controls. A similar requirement for JAK-STAT signaling was observed following *Pe* infection. Control mosquitoes showed robust induction of proliferation, whereas knockdown of JAK or STAT abolished this response, as shown by significantly reduced EdU+ and PH3+ cell counts (Fig. 3d–f). These results demonstrate that JAK-STAT signaling is essential for infection-induced epithelial renewal in *Aedes aegypti* and acts as a conserved regulator of regenerative responses to gut bacterial challenge.

**Figure 3.**
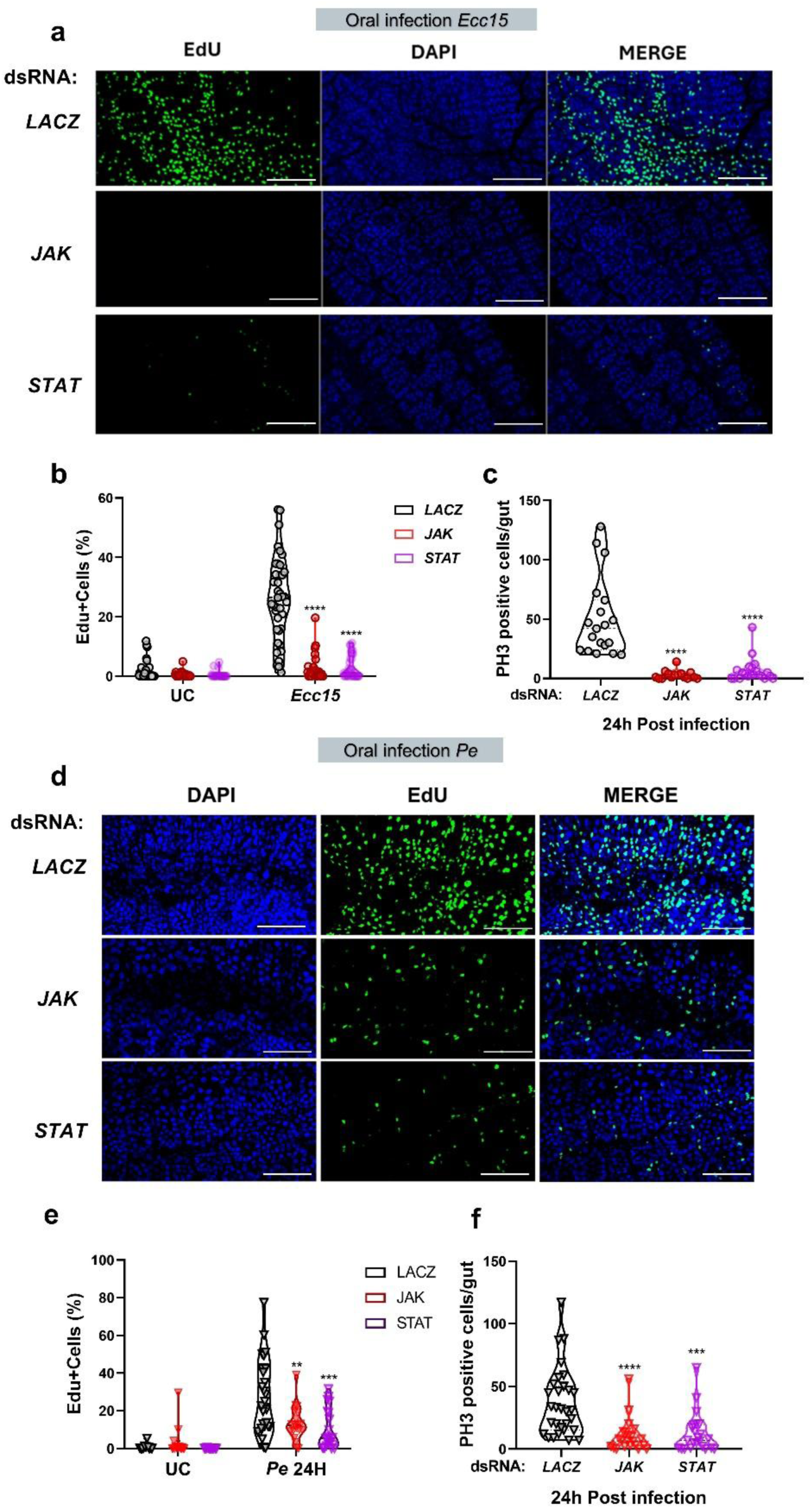
Silencing JAK or STAT impairs intestinal regeneration in response to bacterial infection in *Aedes aegypti*. (a) Representative confocal images of midguts from dsLACZ, dsJAK, and dsSTAT females stained for EdU (green, proliferating cells) and DAPI (blue, nuclei) 24 h after oral infection with *Ecc15*. (b) Quantification of EdU+ cells shows a strong reduction in proliferation in JAK- and STAT-silenced guts compared to controls. (c) Quantification of mitotic cells stained with anti-PH3 antibody also shows reduced mitosis in dsJAK and dsSTAT mosquitoes after *Ecc15* infection. (d) Confocal images showing EdU and DAPI staining in guts 24 h after *Pe* infection. JAK and STAT knockdown prevents proliferation similarly to the *Ecc15* challenge. (e-f) Quantification of EdU+ (e) and PH3+ (f) cells following *Pe* infection confirms that both JAK and STAT are essential for infection-induced regeneration.

### Upd-like is a conserved cytokine-like protein in mosquitoes and shares features with Drosophila Unpaired proteins

We next asked which ligand might activate the JAK-STAT pathway in the mosquito midgut during bacterial infection. Notably, we found that both Vagos (VLG-1 and VLG-2) were not significantly upregulated in the midgut following bacterial challenge (Fig. 2b), suggesting that a different cytokine mediates this response in *Aedes aegypti*. Considering the importance of the JAK-STAT pathway in immunity and now tolerance of infection, we set out to identify the ligand that regulates JAK-STAT upon bacterial infection. We hypothesized that the genome of *Aedes* hosts one or more cytokine-like genes. We further conducted a phylogenetic analysis using amino acid sequences of predicted Unpaired-like proteins across dipteran species. A maximum likelihood tree (Supplementary Fig. 5a) revealed a distinct clade of putative Unpaired-like proteins from culicid mosquitoes, including a previously uncharacterized *Aedes aegypti* protein that we designated Upd-like. This clade is separate from the Drosophila Upd1/2/3 lineage, suggesting that mosquito Upd-like proteins represent an evolutionarily distinct group. These findings support the hypothesis that Upd-like proteins evolved independently in mosquitoes but may fulfill analogous functions to Drosophila Upds in activating JAK-STAT signaling.

To further characterize the *Aedes aegypti* Upd-like protein, we analyzed its primary amino acid sequence for conserved features typical of the Unpaired family (Supplementary Fig. 5b). The N-terminal region contains a predicted signal peptide (boxed in dark green), consistent with secretion. Multiple conserved residues and motifs, particularly proline-, lysine-, and serine-rich regions, were identified (highlighted in red and blue), along with predicted N-linked glycosylation sites and structural elements likely important for receptor binding. These features are indicative of a cytokine-like function and support the hypothesis that *Aedes aegypti* Upd-like may act as a functional analog of Drosophila Unpaired proteins.

To assess evolutionary similarity, we performed pairwise identity and similarity analyses between *Aedes aegypti* Upd-like amino acid sequence and Drosophila Upd1, Upd2, and Upd3 (Fig. 4a). Upd-like showed the highest similarity to Upd2 and Upd3 (∼30%), suggesting potential functional conservation. Interestingly, even Drosophila Upds themselves do not exhibit high similarity or identity values. Structure prediction further revealed intrinsically disordered regions and a central Unpaired-like domain, consistent with cytokine classification (Fig. 4b). We next examined *Upd-like* expression in mosquito midguts following oral bacterial infection. In the Liverpool strain, *Upd-like* was significantly upregulated at 12 hours post-infection with either *Ecc15* (\*\*\*\**p* < 0.0001; Fig. 4c) or *Pe* (\*\**p* < 0.01; Fig. 4d). A similar pattern was observed in the Thai strain, with significant induction following both infections (Fig. 4e), indicating a conserved transcriptional response to enteric bacteria. Altogether, these results identify a putative Unpaired homolog in *Aedes aegypti*, *Upd-like*, that is transcriptionally regulated by enteric bacterial infection and may function as an evolutionarily conserved cytokine activating JAK-STAT signaling.

**Figure 4.**
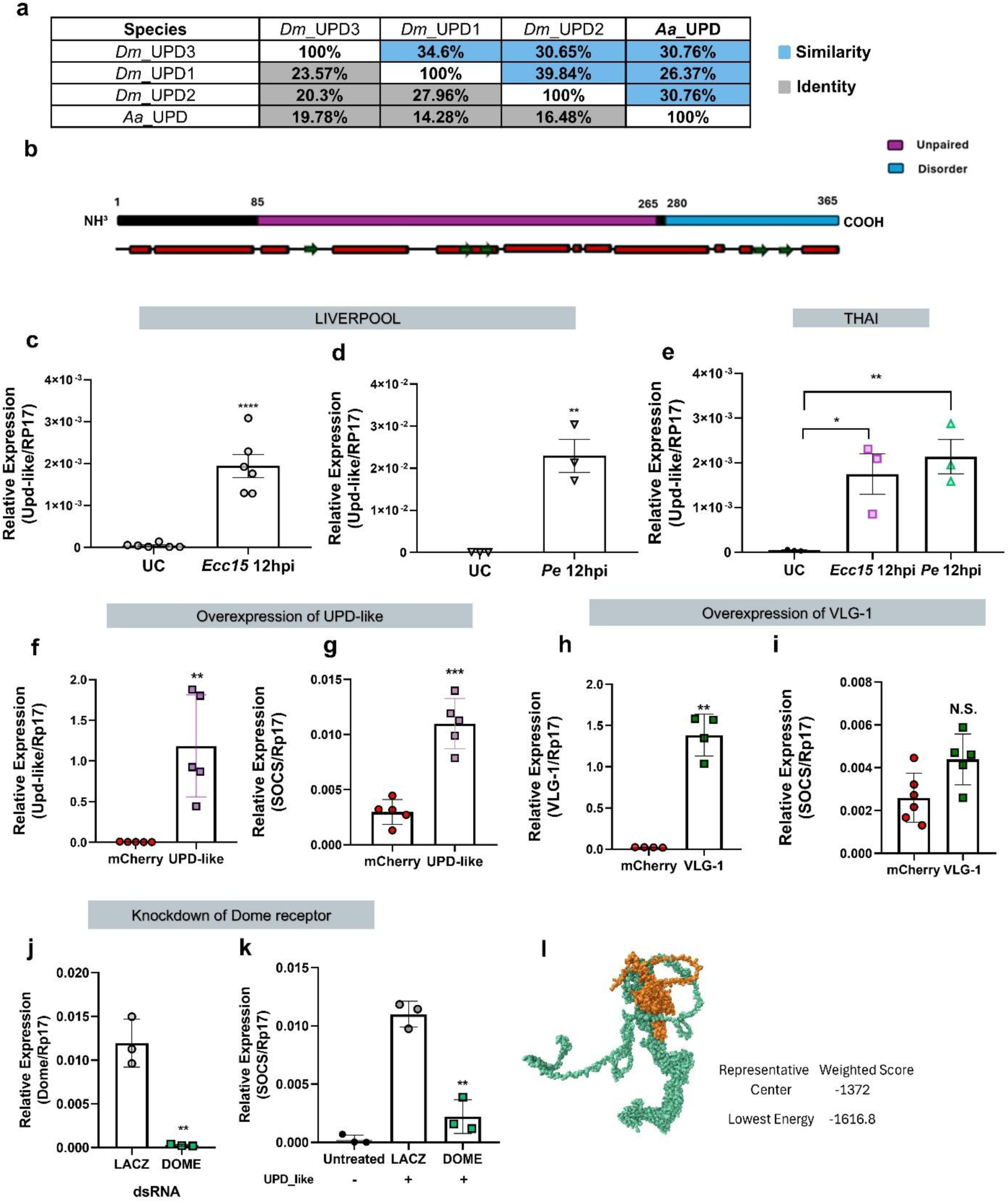
***Upd-like* but not VLG-1 induces JAK/STAT in Aag2 cells.** (a) Sequence identity and similarity between *Drosophila melanogaster* (Dm) Upd ligands and the *Aedes aegypti* (Aa) Upd-like homolog. The *Aedes* protein shows low identity to *Drosophila* ligands but retains structural similarity. (b) Schematic representation of the predicted domain organization of the *Aedes* Upd-like protein, highlighting disordered (blue) and unpaired (purple) regions. (c-e) Upd-like is transcriptionally induced in midguts of Liverpool and Thai *Aedes aegypti* females 12 hours after oral infection with *Ecc15* or PE (p < 0.01–0.0001, t-test). (f-g) Overexpression of Upd-like significantly increases its own expression and induces the JAK-STAT target gene *Socs* (p < 0.01–0.001). (h-i) Overexpression of VLG-1 increases its own transcript but does not induce *Socs*, suggesting pathway specificity for Upd-like in JAK-STAT activation. (j-k) Knockdown of the receptor *Dome* abrogates *Socs* induction by Upd-like overexpression, confirming that Upd-like signals through the JAK-STAT pathway (p < 0.01). (l) Predicted 3D structure of Upd-like (ClusPro 2.0), showing a representative low-energy conformation potentially compatible with receptor binding (lowest energy model: -1616.8).

### *Upd-like* activates the JAK-STAT pathway in Aag2 cells in a Dome-dependent manner

We next asked whether this Upd-like molecule could act as a JAK-STAT ligand. To test this hypothesis, we first performed molecular docking between the predicted structures of Upd-like and the extracellular domain of Dome to assess whether it has the structural potential to act as a ligand for the Domeless receptor. The lowest-energy conformation (ΔG = –1616.8) suggested a stable binding interface (Figure 4l), providing *in silico* structural support for Upd-like functioning as a Dome-dependent cytokine in *Aedes aegypti*.

Next, we tested whether Upd-like functions as a bona fide ligand of the JAK-STAT pathway *in vitro*. In *Aedes aegypti*, two Vago-like genes (VLG-1 and VLG-2) have been identified, neither of which is transcriptionally induced by bacterial infection (Fig. 2b). VLG-1, however, has been reported to possess antiviral activity and is thought to engage the JAK-STAT pathway. To directly compare these candidates, we overexpressed *Upd-like* or VLG-1 in Aag2 cells and measured downstream responses by RT-qPCR. *Upd-like* overexpression significantly increased its own transcript (*p* < 0.01; Fig. 4f) and strongly induced *Socs* expression (*p* < 0.001; Fig. 4g), a canonical target of JAK-STAT signaling. By contrast, VLG-1 overexpression elevated its own transcript (*p* < 0.01; Fig. 4h) but failed to induce *Socs* significantly (Fig. 4i), indicating that, under these conditions, VLG-1 does not activate JAK-STAT. Together, these results identify Upd-like as a functional activator of JAK-STAT signaling in the mosquito midgut.

To determine whether *Upd-like* activates the pathway through the canonical receptor *Domeless* (Dome), we performed RNAi-mediated silencing of Dome in Aag2 cells. Knockdown was efficient (*p* < 0.01; Fig. 4j) and abolished *Upd-like*-induced *Socs* expression (*p* < 0.01; Fig. 4k), indicating that *Upd-like* requires Dome to activate the JAK-STAT pathway. Altogether, these data suggest that *Upd-like* is a bona fide stimulator of the JAK-STAT pathway.

### *Upd-like* Mediates JAK-STAT Activation and Epithelial Renewal in the Midgut

To determine the functional role of *Upd-like* in intestinal homeostasis, we silenced its expression in adult mosquitoes using RNAi (Supplementary Fig. 6a) and examined epithelial responses following enteric bacterial infection. Knockdown of *Upd-like* significantly reduced expression of *Socs*, a canonical JAK-STAT target and negative regulator of the pathway (Supplementary Fig. 6b), confirming that *Upd-like* positively regulates JAK-STAT signaling in the gut. This mirrors findings in Drosophila, where Unpaired (Upd) cytokines activate JAK-STAT and induce *Socs* expression as part of a negative feedback loop to maintain signaling homeostasis^47^. The observed reduction in *Socs* suggests impaired JAK-STAT activity in *Upd-like*–depleted mosquitoes, which may compromise epithelial defense mechanisms during infection.

Following oral infection with *Ecc15*, *Upd-like* knockdown led to a marked reduction in EdU⁺ cells within the midgut epithelium compared to LacZ controls, indicating impaired proliferation (Fig. 5a-b; \*\*\*\**p* < 0.0001). Mitotic activity was similarly diminished, as reflected by a significant decrease in PH3⁺ cells in *Upd-like*–silenced guts (Fig. 5c–d; \*\*\*\**p* < 0.0001). A comparable reduction in proliferation was observed following Pe infection: both EdU incorporation (Fig. 5e; \**p* < 0.05) and PH3⁺ cell counts (Fig. 5f; \*\*\*\**p* < 0.0001) were significantly decreased. These data demonstrate that *Upd-like* acts as a positive regulator of infection-induced epithelial renewal.

**Figure 5.**
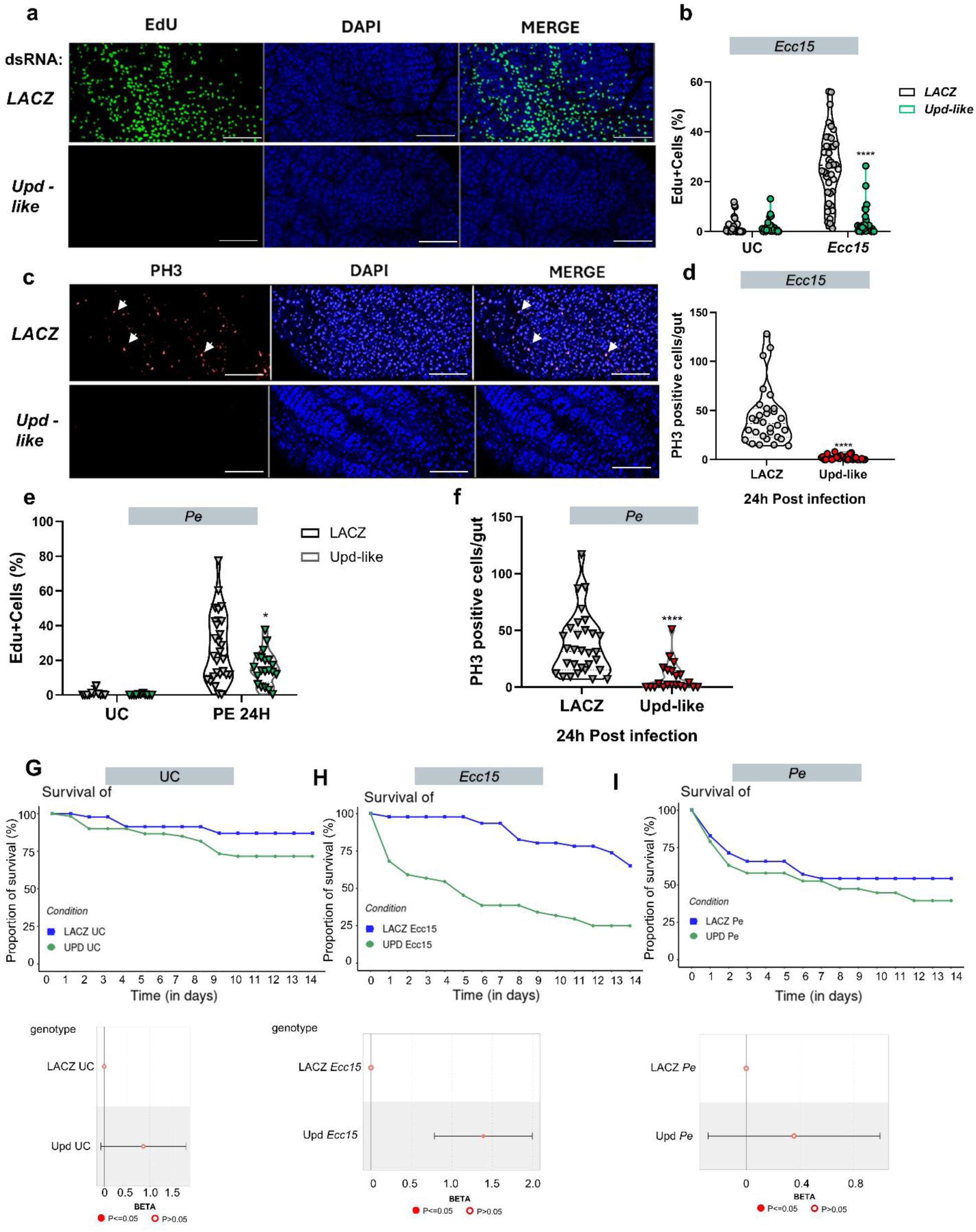
UPD-like knockdown impairs intestinal proliferation and reduces survival after bacterial infection in *Aedes aegypti*. (a) Representative confocal images of midguts stained for EdU (green, proliferation marker) and DAPI (blue, nuclei) from females injected with dsLACZ (control) or dsUPD-like, 24 h after oral infection with *Ecc15*. (b) Quantification of EdU+ cells (%), showing a significant reduction in proliferation in Upd-like-silenced midguts after infection. (c) Representative images of midguts stained with anti-PH3 antibody (red, mitotic marker) and DAPI (blue). (d) Quantification of PH3+ cells per gut at 24 h post-infection with *Ecc15*, showing a strong decrease in mitosis upon Upd-like silencing. (e) EdU+ cell quantification after infection with *Pe*, showing a similar reduction in proliferation. (f) Quantification of PH3+ cells per gut after *Pe* infection, confirming impaired mitosis in Upd-like-silenced mosquitoes. (g-i) Survival curves and statistical analysis of females injected with dsLACZ or dsUPD-like under different conditions: (g) unchallenged (UC), (h) infected with *Ecc15*, and (i) infected with *Pe*. UPD-like knockdown significantly reduces survival only upon bacterial challenge (h and i), suggesting that UPD-like is required for tissue homeostasis during infection. Statistical tests: Mann–Whitney (b, d–f), Log-rank and Cox regression (g–i). \**p* < 0.05; \*\*\*\**p < 0.0001*.

To assess the physiological relevance of this regulation, we measured mosquito survival after infection. *Upd-like* knockdown significantly reduced survival following both *Ecc15* (Fig. 5h) and *Pe* (Fig. 5i) challenge, as shown by Kaplan-Meier curves and Cox model analysis. Importantly, no difference in survival was observed under unchallenged conditions (Fig. 5g), indicating that *Upd-like* is specifically required for infection tolerance rather than basal epithelial maintenance. Together, these findings identify *Upd-like* as a critical activator of the JAK-STAT pathway in the mosquito midgut, promoting epithelial regeneration and host survival during bacterial infection.

## Discussion

Our study identifies the JAK-STAT signaling pathway as a central regulator of epithelial regeneration and host survival in the *Aedes aegypti* midgut following enteric bacterial infection. Rather than serving as a classical immune defense pathway focused on pathogen elimination, JAK-STAT signaling promotes infection tolerance by stimulating intestinal stem cell (ISC) proliferation and epithelial renewal. These findings position the mosquito midgut as a dynamic immune-physiological interface, where immune activation, tissue turnover, and regenerative repair are integrated to maintain homeostasis in the face of diverse microbial challenges.

Mosquito innate immunity is orchestrated by physical barriers, cellular responses (e.g., phagocytosis, melanization), and conserved signaling pathways, including Toll, IMD, JAK-STAT, and RNA interference (RNAi), that collectively detect, control, and tolerate a broad spectrum of pathogens encountered through blood feeding and environmental exposure^21,42,48,49^. In *Drosophila melanogaster*, Toll and IMD activation is strongly pathogen-class specific, with Toll primarily responding to fungi and Gram-positive bacteria, and IMD to Gram-negative bacteria^12^. In contrast, *Aedes aegypti* activates both pathways concurrently in response to enteric bacteria of both Gram types, suggesting a more integrated and convergent immune recognition strategy. This convergence extends beyond the gut, as systemic challenges with diverse bacterial pathogens also trigger a core transcriptional program involving targets from both pathways. Although certain genes such as defensins and cecropins may show preference for particular pathogens, the majority of Imd and Toll pathway targets respond broadly rather than being confined to a single class of infection. This unified activation pattern points to a convergent, multi-pathway immune design in *Aedes aegypti*, revealing consistency between local and systemic responses^34^. This strategy likely reflects evolutionary adaptation to the mosquito’s exposure to highly diverse, microbially rich habitats—providing broad antimicrobial protection while limiting inflammation-mediated tissue damage that could compromise gut integrity^6,7,9,19^.

Within this integrated response, JAK-STAT signaling plays a dual role in *Aedes aegypti*, functioning in both classical antiviral immunity and epithelial regeneration. On one hand, it induces antiviral effectors that restrict viral replication; on the other, it promotes ISC-driven epithelial renewal, preserving barrier function and mitigating pathogen-induced tissue damage. By maintaining epithelial integrity, this regenerative role contributes to infection tolerance and complements pathogen clearance mechanisms. Thus, JAK-STAT signaling coordinates immune activation and tissue repair, two interdependent arms of the host response, underscoring the complexity of mosquito immunity and its impact on vector competence. Striking this balance between resistance and tolerance is essential for mosquito survival in microbe-rich environments^9,26,28,30,42^. However, the precise mechanisms underlying the antiviral functions of JAK-STAT signaling remain poorly understood. We propose that its role in epithelial renewal may itself contribute to both resistance and tolerance, potentially through changes in epithelial composition that influence susceptibility to infection and control of pathogen load.

In this study, we also identify and functionally characterize a previously unrecognized Unpaired (Upd)-like cytokine in *Aedes aegypti*, which is induced upon bacterial infection, activates JAK-STAT signaling, and is essential for ISC proliferation and host survival. While Drosophila encodes three Upd ligands (Upd1–3), each with specialized roles in development, metabolism, and immunity^38,50–52^, mosquitoes appear to rely on a single *Upd-like* gene. This raises key evolutionary questions: Is Upd-like the sole cytokine of this family in mosquitoes? Are additional ligands yet to be discovered? Or did Drosophila expand its Upd family after diverging from the mosquito lineage? Clarifying the evolutionary trajectory of this cytokine family will be critical to understanding how JAK-STAT signaling diversified across insects.

In parallel with Upd-like, the cytokine Vago (VLG-1) has been implicated in activating the JAK-STAT pathway and mosquito antiviral immunity. VLG-1 is induced by viral infections such as West Nile and dengue and acts as a secreted effector that activates JAK-STAT signaling in neighboring cells via a non-canonical, unidentified receptor^41,44,45^. However, our results show that VLG-1 does not activate canonical JAK-STAT signaling in Aag2 cells, as it fails to induce *Socs*, a key transcriptional readout and negative regulator of this pathway^35^. Moreover, although two Vago genes are encoded in the genome, neither is induced by bacterial infection, reinforcing Vago’s role as a virus-specific immune signal. These findings suggest that VLG-1 functions outside the classical JAK-STAT transcriptional loop, potentially engaging distinct mechanisms or receptor systems. The divergence between *Upd-like* and VLG-1 underscores the complexity of mosquito cytokine signaling, where different ligands mediate pathogen-specific responses. Notably, insect Upd family members show structural and functional parallels with mammalian IL-6–like cytokines^38,53^. In mammals, autocrine IL-6 signaling is crucial for maintaining the intestinal stem cell niche and tissue homeostasis, confirming the link between IL-6 activity and epithelial renewal. This highlights the evolutionary conservation of JAK-STAT signaling in coordinating immune defense with tissue repair^54^. Future studies should define the upstream stimuli controlling *Upd-like* expression, its cellular sources and secretion dynamics, and its potential crosstalk with Vago and other effectors; insights that could reveal new layers of mosquito immune regulation and identify novel targets for vector control.

The mosquito midgut epithelium is a highly plastic tissue that responds to developmental, hormonal, microbial, and environmental cues^19,55–57^. ISC-driven epithelial renewal is central to mosquito physiology, supporting digestion, reproduction, lifespan, and resilience to infection^9,56,58^. As in Drosophila, where gut plasticity is modulated by diet, microbiota, and aging^11^, *Aedes aegypti* exhibits comparable dynamic remodeling of its midgut. Recent studies in *Aedes* and *Anopheles* mosquitoes have begun to quantify midgut epithelial dynamics, revealing ongoing ISC proliferation, DNA synthesis, and endoreplication even in adults. These mechanisms allow the midgut to adapt to blood feeding and microbial exposure by increasing epithelial turnover, promoting both mitosis and ploidy elevation to maintain tissue integrity under stress. Notably, the nature and magnitude of these responses vary across species and contexts. For example, in *Anopheles gambiae*, blood feeding primarily induces EC endoreplication rather than progenitor proliferation, reflecting species-specific strategies for maintaining gut function^16^. Additionally, stem cells may also function as immune cells, adding a novel dimension to the gut’s role in both tissue maintenance and immune defense in mosquitoes^48^.These dynamic responses are critical for preserving epithelial homeostasis, shaping microbial tolerance, and influencing vector competence, ultimately supporting the mosquito’s ability to withstand infection and transmit pathogens.

Taken together, our results demonstrate that epithelial renewal in *Aedes aegypti* is driven by a conserved cytokine-receptor module that activates JAK-STAT signaling in response to bacterial infection. This regenerative program promotes host survival by maintaining epithelial integrity and gut function, operating independently of direct pathogen clearance. By placing JAK-STAT at the intersection of immune signaling and epithelial repair, our findings highlight an underappreciated facet of mosquito immunity, one in which sustaining tissue homeostasis, rather than eliminating microbes, emerges as a central strategy for enduring microbial stress and minimizing immunopathology. These insights point to tissue resilience as a critical determinant of vector fitness.

Our findings also suggest that gut regenerative mechanisms may offer novel targets for reducing vector competence. Because epithelial renewal is essential for mosquito survival during infection, interfering with this process, through inhibition of *Upd-like* cytokine production or JAK-STAT activity, could reduce mosquito lifespan or infection tolerance, thereby limiting pathogen transmission. Future studies should focus on (1) identifying downstream effectors of JAK-STAT–dependent regeneration, (2) resolving the spatial and temporal dynamics of immune-regenerative crosstalk, and (3) investigating how chronic or repeated microbial exposure modulates ISC behavior, epithelial composition, and gut barrier function. A deeper understanding of how mosquitoes balance microbial load, tissue damage, and regenerative capacity across life stages and environments will be vital for elucidating the mechanisms of vector resilience and disease transmission.

Ultimately, our work contributes to a growing shift in the field, from a resistance-centric view of vector immunity to an integrated framework that incorporates epithelial plasticity, tissue repair, and infection tolerance as core components of host-pathogen interactions. By uncovering conserved regenerative mechanisms and vector-specific vulnerabilities, we highlight new opportunities for innovative vector control. Future research dissecting mosquito cytokine signaling, stem cell dynamics, and immune– physiological integration will be instrumental in designing strategies to reduce mosquito fitness and transmission potential from within the vector itself.

## Materials and methods

### Mosquito Rearing

*Aedes aegypti* mosquitoes (Liverpool and Thai strains) were reared at Cornell University (Ithaca, NY, USA) under standard insectary conditions. Larvae were maintained at a density of 200 per 1 L of deionized water from the L2 stage to pupation and were fed daily with Hikari fish food (#04428). Adult mosquitoes were maintained on 10% sucrose solution ad libitum. All stages were kept at 29 °C with 75 ± 5% relative humidity and a 12:12 h light/dark cycle.

### Oral Infection

Five-day-old female mosquitoes were starved for 3 h prior to infection. Bacterial challenge was performed by providing mosquitoes with a cotton ball soaked in a 1:1 solution of 20% sucrose and bacterial suspension (OD600 = 2.0 for *Ecc15*; OD600 = 0.1 for *P. entomophila*). Mosquitoes were allowed to feed for 4 h, after which they were returned to 10% sucrose.

### RNA Extraction and Sequencing

For RNA-seq, midguts from 20 mosquitoes per condition were dissected at 4, 12, and 24 h post-infection (three biological replicates per condition), homogenized in 500 μL TRIzol (Invitrogen), and stored at −80 °C. RNA was extracted using a phenol-chloroform protocol. Libraries were prepared using the Lexogen QuantSeq 3′ mRNA-Seq kit and sequenced as 75 bp single-end reads on an Illumina NextSeq 500 at the Cornell Life Sciences Core. RNA quality and concentration were verified by spectrophotometry and sample quality was evaluated before and after library preparation using a fragment analyzer (Advanced Analytical). Raw high throughput sequencing data are deposited to SRA repository under accession number: SRA: PRJNA1331784.

### RNAseq data analysis

Read quality was assessed with FastQC (https://github.com/s-andrews/FastQC). Reads were trimmed using BBMap (https://jgi.doe.gov/data-and-tools/bbtools/), then aligned to the Ae. aegypti LVP_AGWG AaegL5.2 transcriptome (VectorBase) using Salmon v0.9.1. Transcript-level counts were summed per gene and analyzed with DESeq2. Principal component analysis (PCA) was performed in R using custom scripts. GO enrichment and InterPro domain analyses were conducted with topGO using the classic Fisher method.

### Statistical analysis

Experiments included ≥3 biological replicates unless stated otherwise. For qPCR, two-tailed unpaired Student’s t-tests were used. Survival data were analyzed via the Mantel– Cox log-rank test. All data are presented as mean ± SE. P values <0.05 were considered statistically significant. Statistical details are provided in figure legends.

### cDNA synthesis and qPCR

Total RNA was extracted from dissected midguts or Aag2 cells and reverse transcribed using the qScript cDNA Synthesis Kit (Quantabio). qPCR reactions were performed with PerfeCTa SYBR Green SuperMix (Quantabio) in 15 μL volumes using gene-specific primers (Supplementary Table 1). Reactions were run in triplicate, and melting curve analysis confirmed amplicon specificity. Relative gene expression was calculated by the 2^−ΔΔCt method, normalized to RPS17.

### dsRNA Synthesis and Knockdown

Double-stranded RNAs (dsRNAs) targeting *Domeless*, JAK, STAT, and *Upd-like* were synthesized using the MEGAscript T7 Kit (Invitrogen) according to manufacturor’s instructions For in vivo knockdown, ∼0.5 μg of purified dsRNA was injected intrathoracically into cold-anesthetized, 5-day-old females. Control mosquitoes received dsLacZ. For *in vitro* silencing, Aag2 cells were transfected with 1 μg dsRNA/well. Knockdown efficiency was validated by RT-qPCR at 12 h (mosquitoes) or 48 h (cells) post-treatment.

### Proliferation Assay (EdU Incorporation)

Proliferation was measured by EdU incorporation. EdU (200 μM; Click-iT™ EdU Cell Proliferation Kit, ThermoFisher) was added to the sucrose-bacteria mixture during feeding. Midguts were dissected at 24 h post-infection, fixed in 4% paraformaldehyde, permeabilized in 0.15% Triton X-100, and processed per manufacturer’s instructions. Nuclei were counterstained with DAPI (1 μg/mL), and samples were mounted in Citifluor AF1. Images were acquired using a Zeiss LSM 700 confocal microscope.

### Quantification of mitotic cells (PH3 Staining)

Midguts were dissected in PBS, fixed in 4% paraformaldehyde (30 min, RT), permeabilized in 0.15% Triton X-100, and blocked in PBS with 0.1% Tween-20, 2.5% BSA, and 10% normal donkey serum. Tissues were incubated with rabbit anti-PH3 antibody (1:500, Merck Millipore), followed by Alexa Fluor 555-conjugated donkey anti-rabbit IgG (1:2000, ThermoFisher). Guts were mounted and imaged as above.

### Survival Assays

Liverpool mosquitoes were injected with dsRNA and allowed to recover for 48 h before bacterial challenge. Twenty mosquitoes per condition were monitored daily for up to 14 days. Survival curves were analyzed using the log-rank (Mantel–Cox) test.

### Bacterial load Quantification (CFU assay)

At 6, 12, 24, 48, and 72 h post-infection, individual mosquitoes (n = 10 per condition) were homogenized in 500 μL sterile PBS. Homogenates were incubated overnight at 4 °C, serially diluted (1:100, 1:1000), and plated on LB agar with rifamycin (50 μg/mL). Plates were incubated at 29 °C for 24 h, and colony counts were recorded. Each condition was performed in triplicate.

### Protein-Protein Docking Prediction

Interactions between Upd-like and the extracellular domain of Domeless were predicted using the ClusPro web server. PDB-formatted structures were uploaded, and ClusPro generated low-energy binding models based on rigid-body docking and clustering of top 1000 orientations^59,60^.

### Phylogenetic Analysis, similarity and Identity analysis

Putative Unpaired-like protein sequences from dipteran species were identified using PSI-BLAST (Position-Specific Iterated BLAST) searches against the NCBI non-redundant protein database, using the *Aedes aegypti* Upd-like sequence as the query. PSI-BLAST was run with default parameters for three iterations to ensure sensitive detection of both close and distant homologs. All retrieved sequences were aligned using MUSCLE as implemented in MEGA X. The resulting multiple sequence alignment was manually inspected and trimmed to remove poorly aligned or divergent regions. Phylogenetic trees were constructed in MEGA X.

To assess the degree of similarity and identity between the Unpaired (Upd) protein from *Drosophila melanogaster* and the Upd-like protein from *Aedes aegypti*, we performed pairwise sequence comparisons using the Sequence Identity and Similarity (SIAS) server. The amino acid sequences for Drosophila Upd (Upd1, Upd2, and Upd3) were retrieved from the NCBI database, and the predicted Upd-like sequence from *Aedes aegypti* was obtained from VectorBase. The sequences were submitted to the SIAS (Sequence Identity and Similarity) web server (http://imed.med.ucm.es/Tools/sias.html), which calculates both percentage identity and similarity based on the BLOSUM62 substitution matrix. Pairwise alignments were performed using default parameters, and the resulting values were recorded for each comparison. The SIAS output provided a quantitative measure of evolutionary conservation and potential functional similarity between the mosquito Upd-like and Drosophila Unpaired proteins

### Overexpression of *Upd-like* and VLG-1

Coding sequences of *Upd-like* and VLG-1 were cloned into a modified pFastBac™ (pFBactin-linker) under the Drosophila *Actin5C* promoter. The control NLS-mCherry bacmid was generated by BamHI cloning the mCherry ORF (in place of the EGFP in FBactin:NLS-EGFP^61^, fused in-frame with a C-terminal NLS sequence. All pFastBac-derived transfer vectors were introduced into DH10Bac™ *E. coli* for bacmid generation. Recombinant bacmid DNA was transfected into Sf9 cells (grown in Sf900III medium) to produce viral particles, which were then amplified in Sf9 cells, titrated by end-point dilution in Sf9-ET cells^62^. Virus particles were used to transduce Aag2 cells (grown in Sf900III medium) using three infectious units per cell, then and incubated at 27°C for 24 h before collecting cells for analysis of gene expression. Overexpression of VLG-1 and *Upd-like* was confirmed by RT-qPCR.

### Cell Transfection

Aag2 cells grown in Schneider’s medium (+8% FBS) were seeded at 2 × 10⁵ cells/well in 24-well plates and incubated overnight. For transfection, 1 μg of dsRNA was combined with Attractene Reagent (Qiagen) in 150 μL serum-free Schneider’s medium, incubated 30 min, and applied to cells. Medium was topped to 300 μL, and after 1 h at 28 °C, 300 ul of growth medium containing 16% FBS was added. Cells were harvested at 48 h post-transfection for RT-qPCR.

## Funding

This research was supported by grants from NIH (R01AI148529 and R01AI148541 to NB) and NSF (IOS 2024252 to NB).

## Acknowledgments

We would like to thank the members of the Buchon lab Peter Nagy, George-Rafael Samantsidis, Robin Chen and Xuerong Jin for helpful discussions, suggestions and comments on the manuscript. A special thank you to Laura Harrington for her kind provision of Thai strain mosquito.

## Supplementary Figures

**Supplementary Figure 1.**
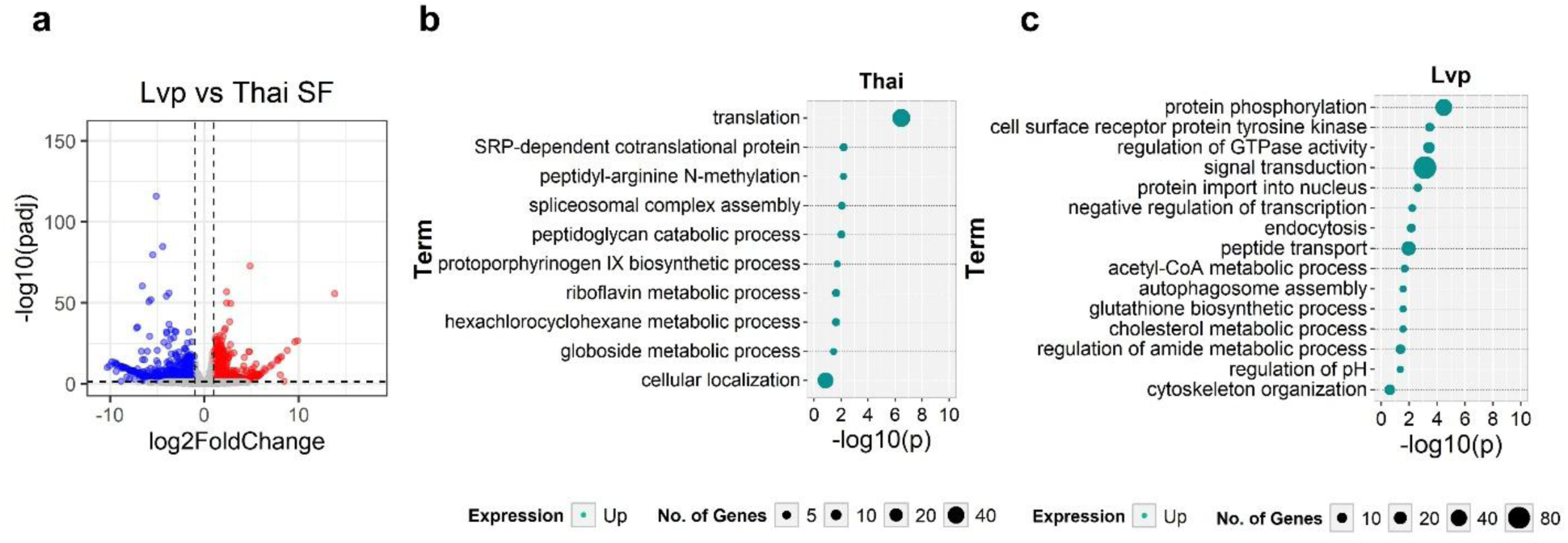
Transcriptomic comparison between Thai and Liverpool (Lvp) strains under unchallenged conditions. (a) Volcano plot showing differentially expressed genes between Lvp and Thai strains (Lvp/Thai). Red dots represent genes significantly upregulated in Lvp, and blue dots represent genes significantly upregulated in Thai. (b) Gene Ontology (GO) enrichment analysis of genes higher expressed in Thai, showing significantly enriched biological processes. (c) GO enrichment analysis of genes higher expressed in Lvp, showing significantly enriched biological processes. Circle size reflects the number of genes associated with each GO term, and the x-axis indicates enrichment significance (−log10 *p*).

**Supplementary Figure 2.**
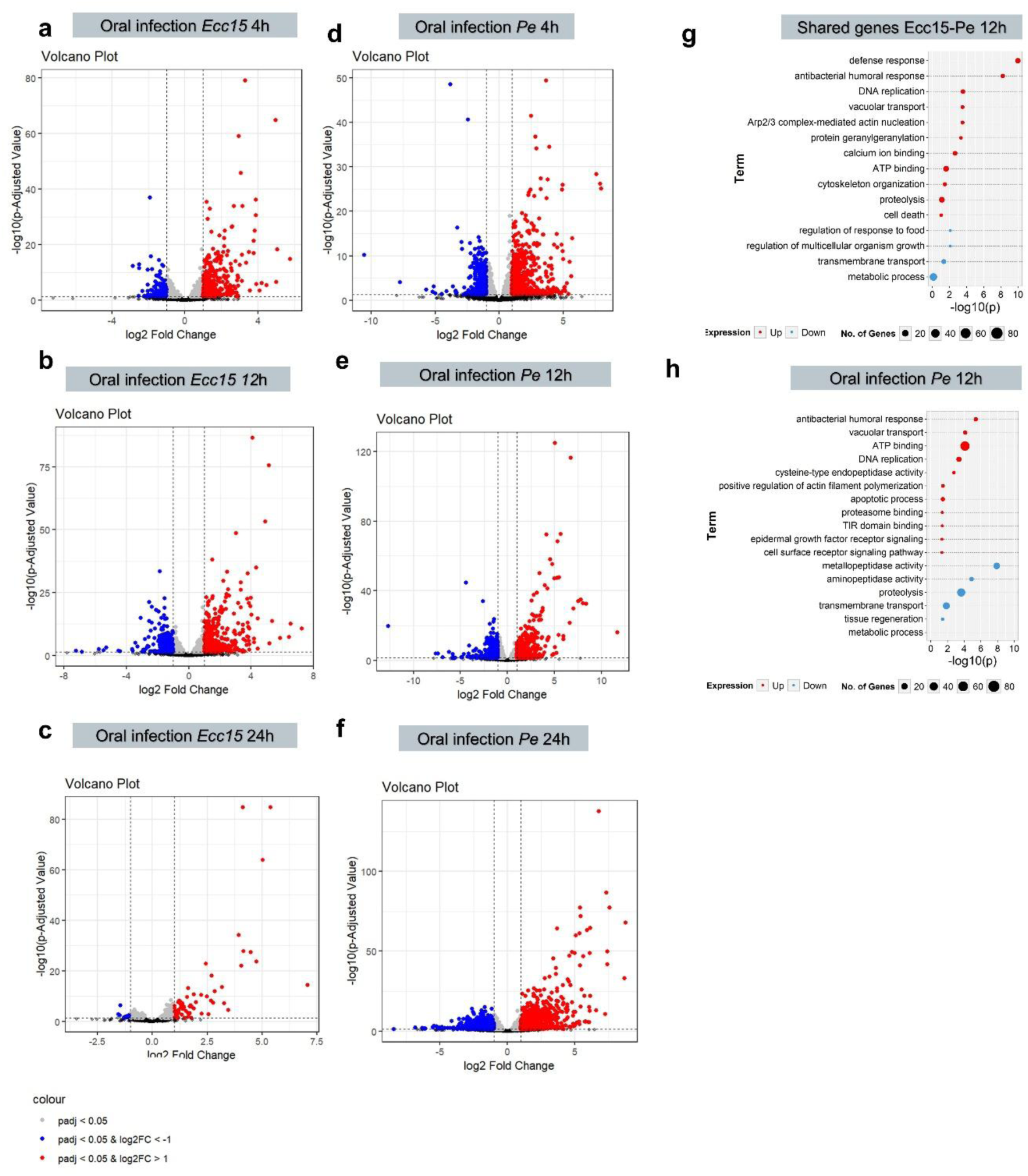
Transcriptional response of mosquito midgut to bacterial infection. (a–f) Volcano plots showing differential gene expression in mosquito midguts following oral infection with *Ecc15* (a–c) and *Pe* (d–f) at 4h (a, d), 12h (b, e) and 24h (c, f) post-infection. The x-axis represents log2 fold change, and the y-axis represents –log10 of the adjusted p-value (padj). Red points indicate significantly upregulated genes (padj < 0.05 and log2 fold change > 1), blue points indicate significantly downregulated genes (padj < 0.05 and log2 fold change < –1), and grey points indicate genes without significant differential expression. (g) Comparative GO enrichment analysis of differentially expressed genes between *Ecc15* and *Pe* at 12h post-oral infection. (h) GO enrichment plot for differentially expressed genes at 12h post-infection with *Pe*. GO terms (y-axis) are plotted against –log10(padj) (x-axis). Circle size indicates the number of genes in each term; circle color shows the direction of gene expression (red: upregulated, blue: downregulated).

**Supplementary Figure 3.**
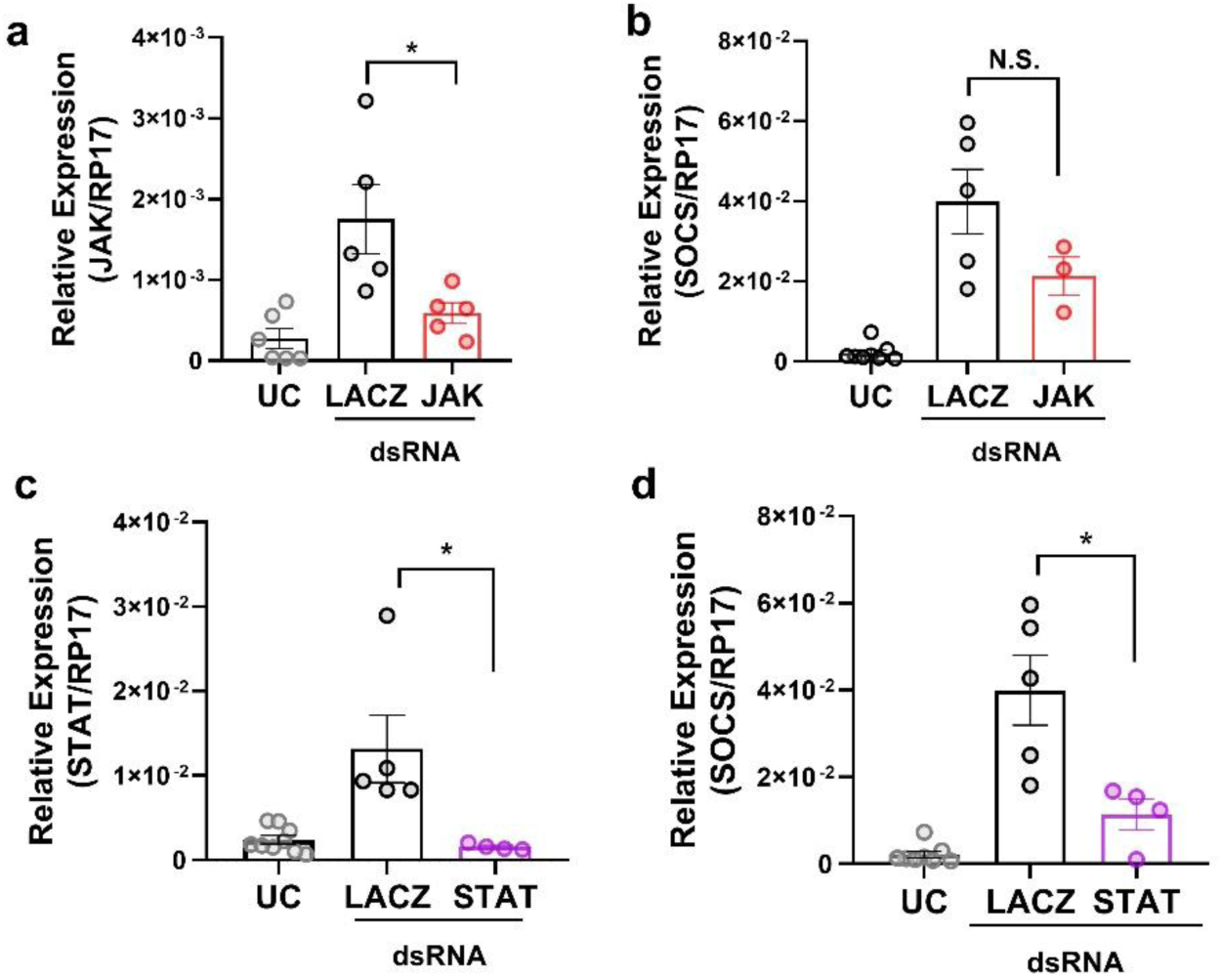
JAK-STAT pathway knockdown reduces *Socs* expression in *Aedes aegypti*. Quantitative RT-PCR analysis of gene expression in midgut tissues from adult female Aedes aegypti mosquitoes injected with dsRNA targeting either LacZ (control), JAK, or STAT, and analyzed 12 hours post-infection.(a) Knockdown efficiency of JAK relative to control (LacZ) and untreated controls (UC).(b) Expression of *Socs* following JAK knockdown.(c) Knockdown efficiency of STAT.(d) Expression of *Socs* following STAT knockdown. Gene expression is normalized to the housekeeping gene RPS17. Bars represent mean ± SE, and individual data points represent biological replicates (n = 20 per replicate). *indicates statistically significant difference compared to LacZ control (*p* < 0.05, determined by t-test).

**Supplementary Figure 4.**
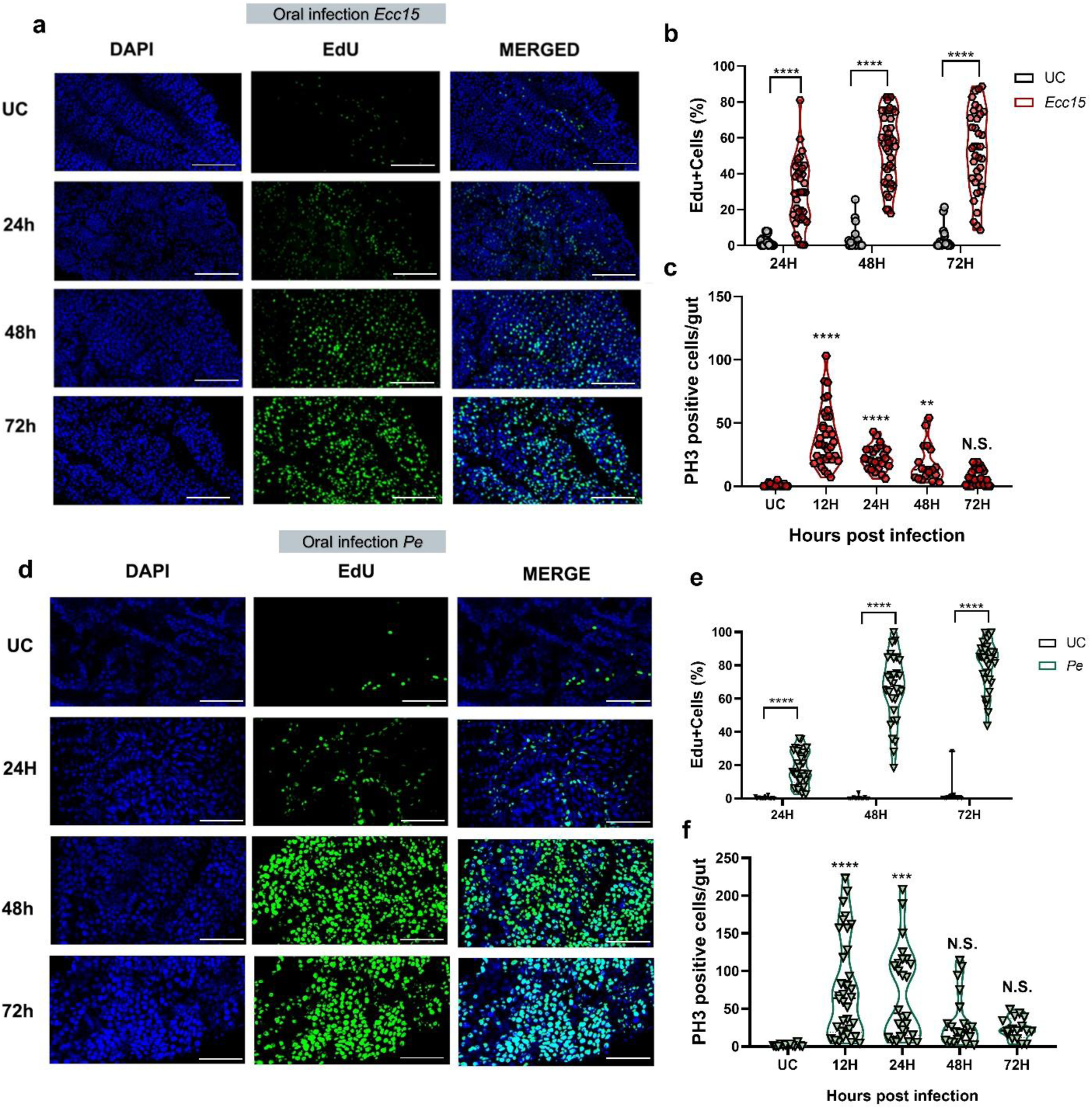
Intestinal epithelial proliferation increases following oral bacterial infection. (a-c) Representative confocal images (a) and quantification of EdU+ cells (b) and PH3+ cells (c) in midguts from mosquitoes infected orally with *Ecc15* at 24, 48, and 72 hours post infection (hpi). EdU incorporation (green) marks cells in S-phase, while PH3 staining identifies mitotic cells. Nuclei are counterstained with DAPI (blue). Compared to unchallenged (UC) controls, *Ecc15*-infected guts show a robust increase in EdU+ cells from 24 hpi onward and a transient spike in mitotic figures at 12 hpi, suggesting early activation of proliferation followed by sustained DNA synthesis. (d–f) Similar analysis of posterior midguts from mosquitoes infected with *P. entomophila* (Pe), with representative images (d) and quantification of EdU+ (e) and PH3+ (f) cells. As with *Ecc15*, Pe infection induces an early increase in mitotic activity at 12 hpi and elevated DNA synthesis from 24 to 72 hpi, consistent with activation of epithelial renewal programs.

**Supplementary Figure 5.**
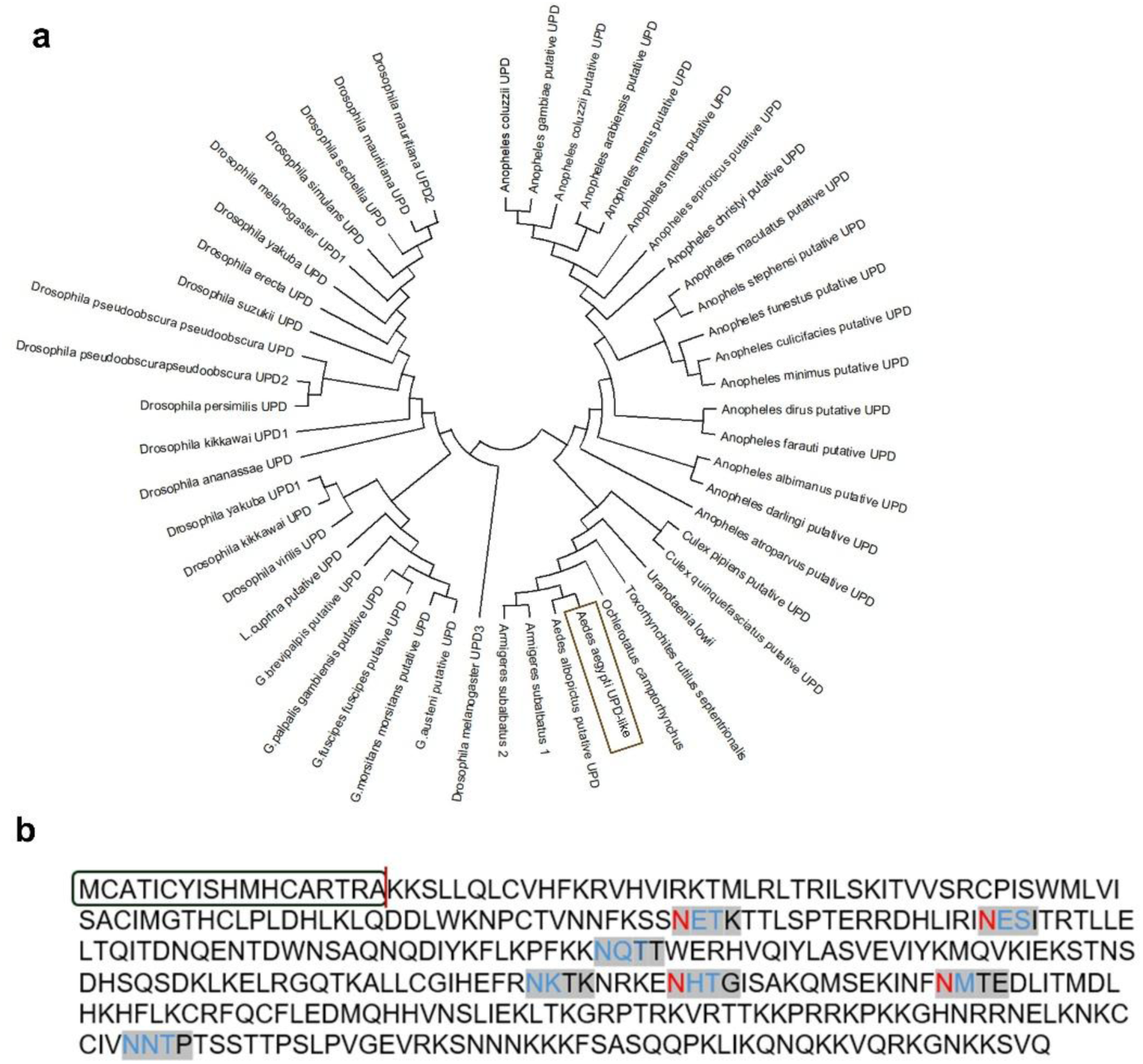
Sequence conservation and structural features of a putative *Aedes aegypti* UPD-like protein. (a) Phylogenetic analysis of UPD-like proteins from various dipteran species, including multiple *Drosophila* and mosquito species. The tree highlights a distinct clade of UPD proteins in *Drosophila* and a broader group of putative homologs in mosquitoes, indicating potential evolutionary conservation of function despite sequence divergence. (b) Amino acid sequence of the putative *Aedes aegypti* UPD protein. Putative N-glycosylation motifs (Asn-Xaa-Ser/Thr sequons) are highlighted in **blue**, and asparagine residues within those sequons that are predicted to be N-glycosylated are shown in **red**. A signal peptide is boxed (dark green), indicating secretion potential. The presence of glycosylation motifs supports the hypothesis that this protein may undergo post-translational modification typical of extracellular ligands.

**Supplementary Figure 6.**
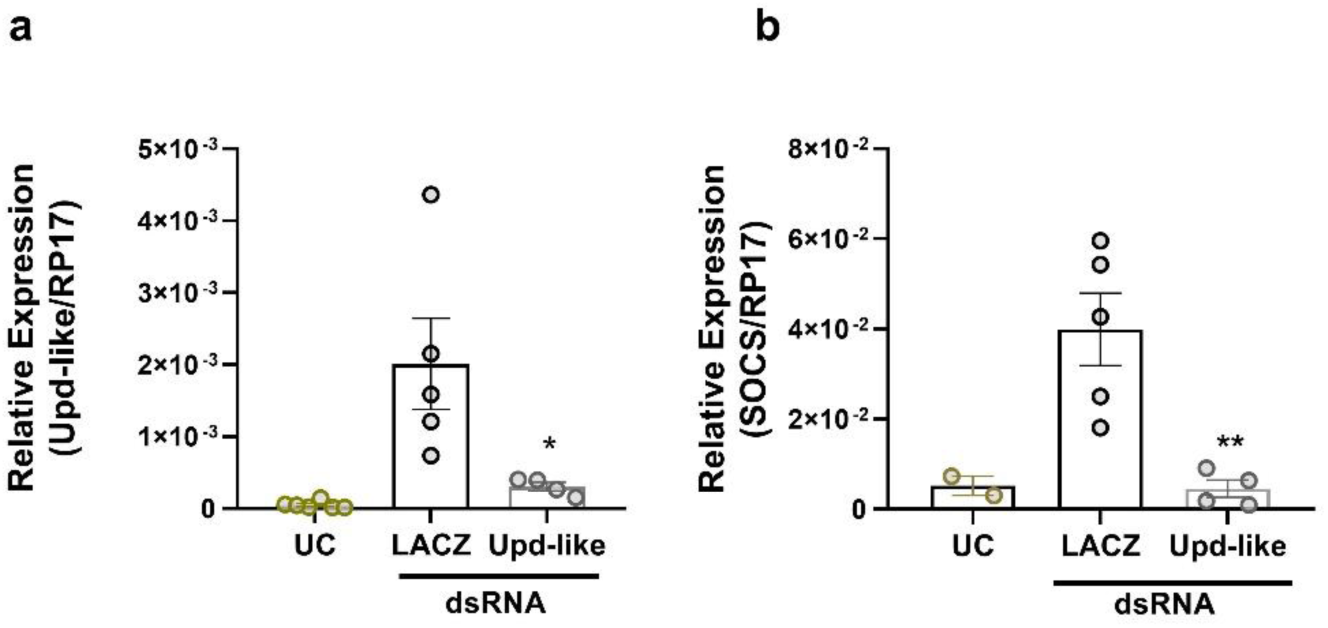
UPD-like knockdown reduces *Socs* expression in the *Aedes aegypti* midgut. RNAi-mediated silencing of *UPD-like* effectively reduces its own transcript levels and leads to a marked decrease in *Socs* expression, indicating that UPD-like positively regulates *Socs* and thus JAK-STAT pathway activity in the mosquito midgut. (a) Relative expression of *UPD-like* (normalized to RP17) in midguts of unchallenged (UC) mosquitoes and those injected with dsRNA targeting LacZ (control) or *UPD-like*. *UPD-like* transcript levels are significantly reduced in the *UPD-like* dsRNA group compared to LacZ controls (*p < 0.05). (b) Relative expression of *Socs* (normalized to RP17) in the same groups. *Socs* expression is significantly decreased following UPD-like knockdown compared to LacZ controls (**p < 0.01). Bars represent mean ± SEM; individual data points are shown.

**Supplementary table 1.**
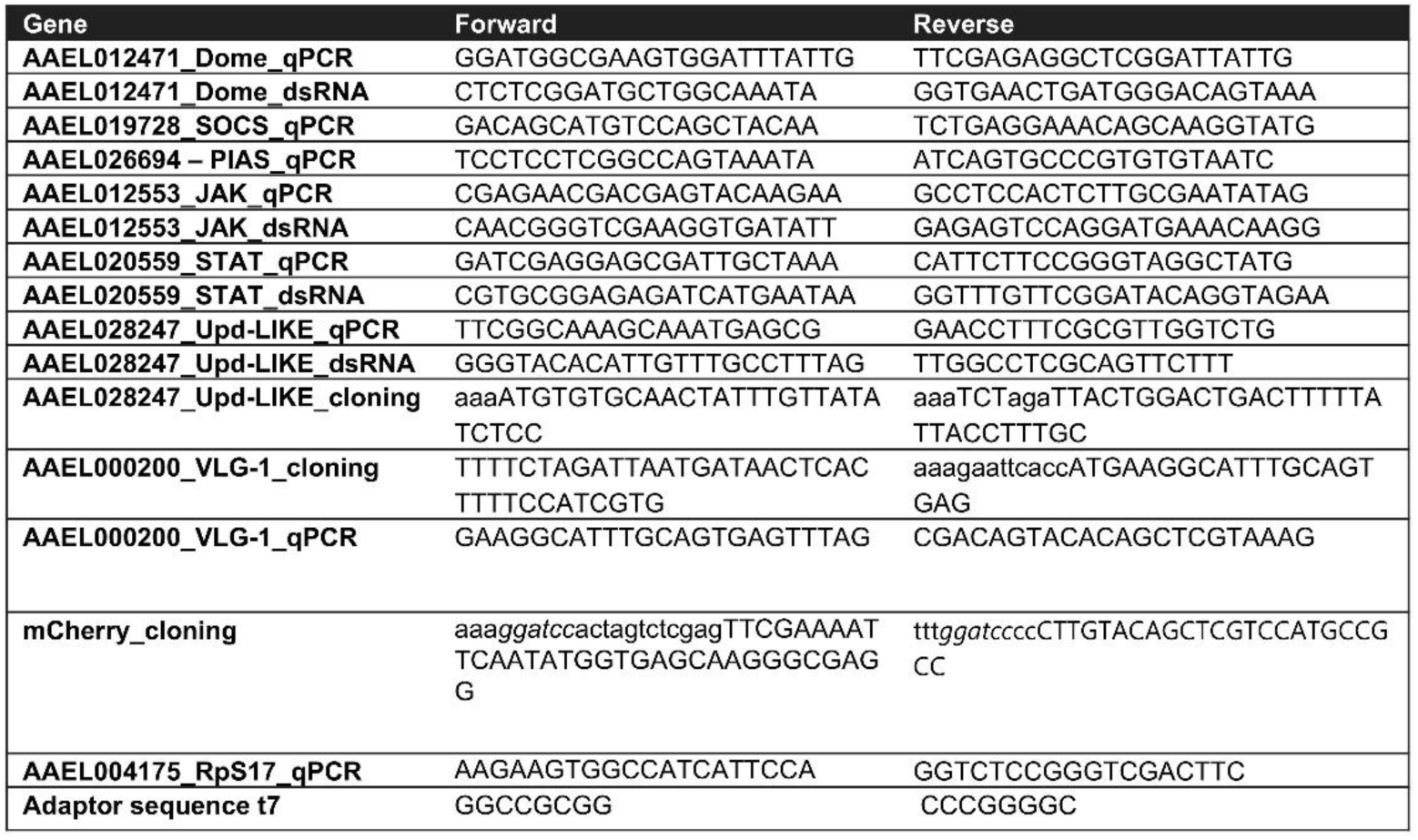
Primers sequences used for qpcr, dsrna synthesis and cloning.

## Notes

### Competing Interest Statement

The authors have declared no competing interest.

